# Fully Automated Brain Tumor Segmentation and Survival Prediction of Gliomas using Deep Learning and MRI

**DOI:** 10.1101/760157

**Authors:** Chandan Ganesh Bangalore Yogananda, Sahil S. Nalawade, Gowtham K. Murugesan, Ben Wagner, Marco C. Pinho, Baowei Fei, Ananth J. Madhuranthakam, Joseph A. Maldjian

## Abstract

Tumor segmentation of magnetic resonance (MR) images is a critical step in providing objective measures of predicting aggressiveness and response to therapy in gliomas. It has valuable applications in diagnosis, monitoring, and treatment planning of brain tumors. The purpose of this work was to develop a fully automated deep learning method for brain tumor segmentation and survival prediction. Well curated brain tumor cases with multi-parametric MR Images from the BraTS2019 dataset were used. A three-group framework was implemented, with each group consisting of three 3D-Dense-UNets to segment whole tumor (WT), tumor core (TC) and enhancing tumor (ET). This method was implemented to decompose the complex multi-class segmentation problem into individual binary segmentation problems for each sub-component. Each group was trained using different approaches and loss functions. The output segmentations of a particular label from their respective networks from the 3 groups were ensembled and post-processed. For survival analysis, a linear regression model based on imaging texture features and wavelet texture features extracted from each of the segmented components was implemented. The networks were tested on the BraTS2019 validation dataset including 125 cases for the brain tumor segmentation task and 29 cases for the survival prediction task. The segmentation networks achieved average dice scores of 0.901, 0.844 and 0.801 for WT, TC and ET respectively. The survival prediction network achieved an accuracy score of 0.55 and mean squared error (MSE) of 119244. This method could be implemented as a robust tool to assist clinicians in primary brain tumor management and follow-up.

## 1. INTRODUCTION

Gliomas account for the most common malignant primary brain tumors in both pediatric and adult populations [1]. They arise from glial cells and are divided into low grade and high grade gliomas with significant differences in patient survival. Patients with aggressive high grade gliomas have life expectancies of less than 2 years [2]. Glioblastoma (GBM) are aggressive brain tumors classified by the world health organization (WHO) as stage IV brain cancer [3, 4]. The overall survival for GBM patient is poor and is in the range of 12 to 15 months [5–7]. These tumors are typically treated by surgery, followed by radiotherapy and chemotherapy. Gliomas often consist of active tumor tissue, necrotic tissue and surrounding edema. Magnetic Resonance Imaging (MRI) is the most commonly used modality to assess brain tumors because of its superior soft tissue contrast. It is routinely used in the clinical work-up of patients for brain tumor diagnosis, monitoring progression and treatment planning [8, 9]. Each MR sequence provides specific information about different tissue sub-components of gliomas. For instance, T1-weighted images with intravenous contrast highlight the most vascular regions of the tumor, called ‘enhancing tumor’ (ET), along with the ‘tumor core’ (TC) that does not involve peri-tumoral edema. Conventional T2-weighted (T2W) and T2W-Fluid Attenuation Inversion Recovery (FLAIR) images are used to evaluate the tumor and peri-tumoral edema together defined as the ‘whole tumor’ (WT) [10].

MRI tumor segmentation is used to identify the subcomponents as enhancing, necrotic or edematous tissue. Due to heterogeneity and tissue relaxation differences in these subcomponents, multi-parametric (or multi-contrast) MRI are often used simultaneously for accurate segmentation [11]. Manual brain tumor segmentation is a challenging and tedious task for human experts due to the variability of tumor appearance, unclear borders of the tumor and the need to evaluate multiple MR images with different contrasts simultaneously [12]. In addition, manual segmentation is often prone to significant intra- and inter-rater variability [12, 13]. Hence machine learning algorithms have been developed for tumor segmentation with high reproducibility and efficiency [12–14]. Following the early success of CNNs [14, 15], they are used as one of the major machine learning methods to achieve great success in clinical applications. [16, 17]. Furthermore, underlying molecular heterogeneity in gliomas makes it difficult to predict the overall survival (OS) of GBM patients based on MR imaging alone [16, 17]. Clinical features [5], along with MR imaging based texture features [18–20] have been used to predict the OS in GBM patients. In this work, we utilized designed a 3D Dense-Unet for segmenting brain tumors into subcomponents and used MRI based texture features from each of these subcomponents for survival prediction in GBM patients. The purpose of this work was to develop a deep learning method with high prediction accuracy for brain tumor segmentation and survival prediction that can be easily incorporated into the clinical workflow.

## 2. MATERIAL & METHODS

### 2.1 Data & Pre-Processing

#### 2.1.1 Brain Tumor Segmentation

335 well curated multi-parametric brain MR images including T2w, T2w-FLAIR, T1w and T1C (post contrast) from the BraTS2019 dataset were used [10, 21–24]. The dataset consisted of 259 high grade glioma (HGG) cases and 76 low grade glioma (LGG) cases. The dataset also included three ground truth labels for a) enhancing tumor, b) non-enhancing tumor including necrosis and c) edema. 125 cases from the BraTS2019 validation dataset was used for evaluating the network’s performance. Pre-processing steps included N4BiasCorrection to remove the RF inhomogeneity [25] and intensity normalization to zero-mean and unit variance.

#### 2.1.2 Survival Analysis

259 HGG subjects were provided for the BraTS2019 Survival challenge. Information regarding age, survival days and resection status (Gross Total Resection (GTR), Subtotal Resection (STR) or not available (NA)) were also provided. 210 subjects out of 259 subjects were selected, as the other 49 subjects were either alive or survival days were not available (NA). The results were evaluated on 29 GTR subjects from the validation dataset. Pre-processing steps were similar to the segmentation task including N4BiasCorrection and intensity normalization.

### 2.2 Network Architecture

Brain tumors contain a complex structure of sub-components and are challenging for automated tumor segmentation. Specifically, the appearance of enhancing tumor and non-enhancing tumor are often different between HGG and LGG. In order to simplify the complex segmentation problem, first a simple convolutional neural network (CNN) was developed to classify HGG and LGG cases. Next, to maintain consistency with the output results provided by the BraTS challenge, separate networks were trained to recognize and predict whole tumor (consisting of enhancing + non-enhancing + necrosis + edema; Whole-net), tumor core (consisting of enhancing + non-enhancing + necrosis; Core-net), and enhancing tumor (enhance-net) as binary features using 3D Dense UNets. Each of these three networks were designed separately for HGG and LGG cases.

The networks were designed to predict the local structures for tumors and sub-components in multi-parametric brain MR images. The architecture of each network is shown in Fig 1. On the encoder side of the network, multi-parametric images passed through initial convolution to generate 64 feature maps to be used in subsequent dense blocks. Each dense block consisted of five layers as shown in Fig 1. Each layer included four sublayers, BatchNormalization, Rectified Linear Unit (ReLu), 3D Convolution and 3D Spatial dropout that were connected sequentially. At each layer, the input was used to generate k feature maps (referred to as growth rate and set to 16) that were subsequently concatenated to the next input layer. The next layer was then applied to create another k feature maps. To generate the final dense block output, inputs from each layer were concatenated with the output of the last layer. At the end of each dense block, the input to the dense block was also concatenated to the output of that dense block. The output of each dense block followed a skip connection to the adjacent decoder. In addition, each dense block output went through a transition down block until the bottle neck block. With this connecting pattern, all feature maps were reused such that every layer in the architecture received a direct supervision signal [26]. On the decoder side, a transition up block preceded each dense block until the final convolution layer followed by a sigmoid activation layer. Two techniques were used to circumvent the problem of maintaining a high number of convolution layers. A) The bottle neck block (Dense block 4 in figure 1 was used such that if the total number of feature maps from a layer exceeded the initial number of convolution maps (i.e. 64) then it was reduced to 1/4^th^ of the total generated feature maps in that layer. B) A compression factor of 0.75 was used to reduce the total number of feature maps after every block in the architecture. In addition, due to the large number of high resolution feature maps, a patch based 3D Dense-Unet approach was implemented where higher resolution information was passed through the standard skip connections.

**Figure 1:**
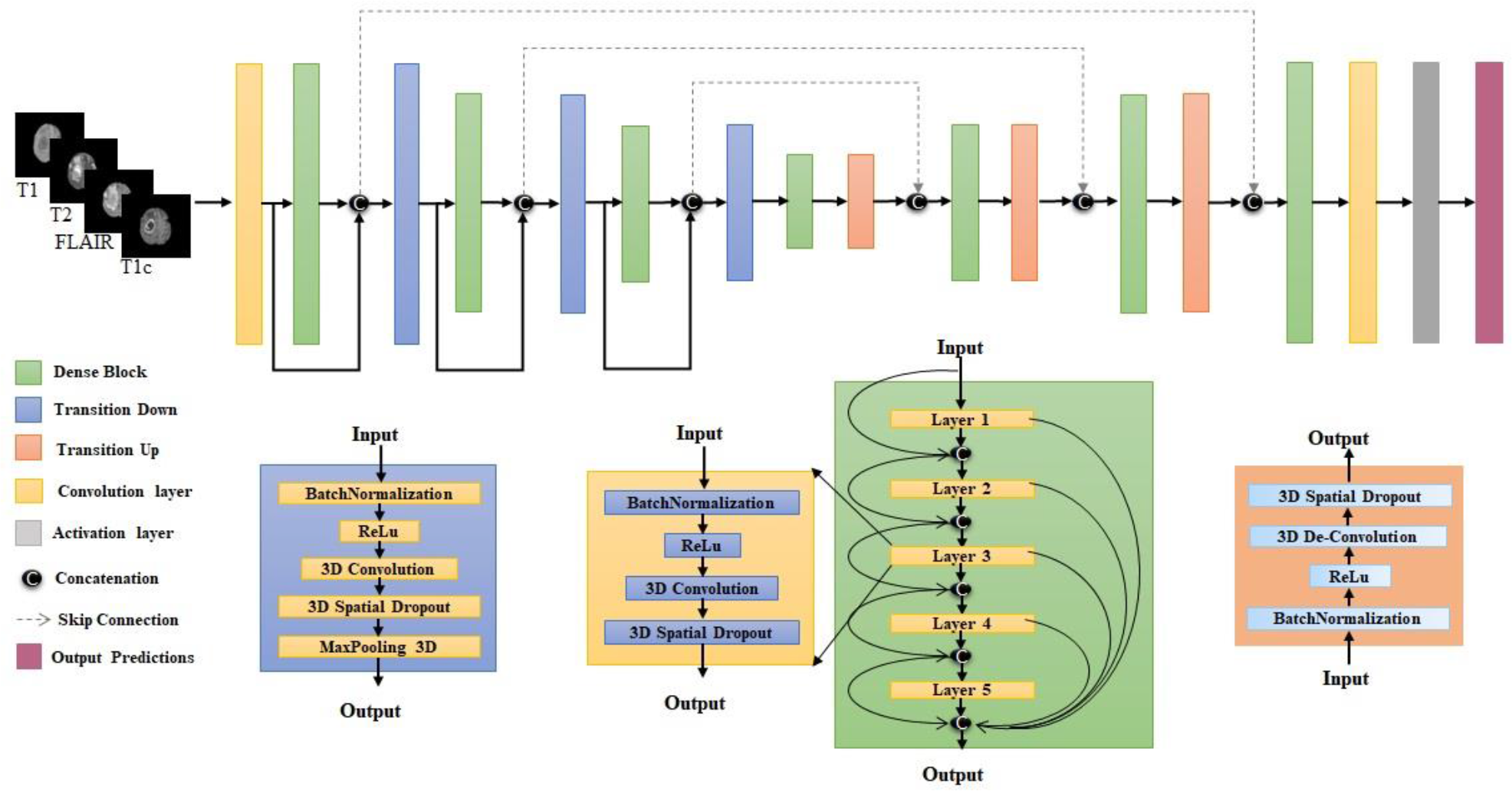
Network Architecture

**Figure 2:**
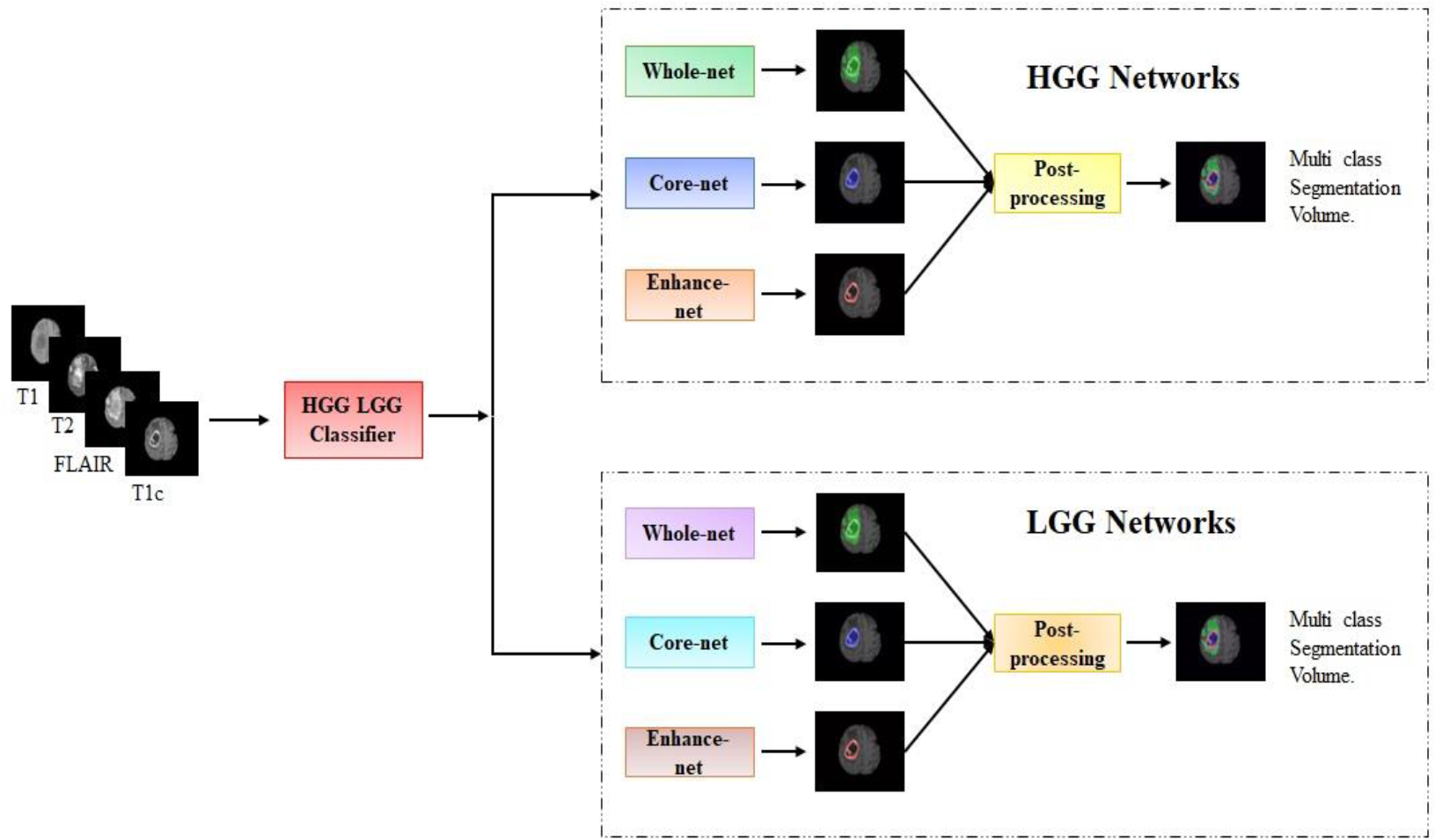
Overview of the 3rd approach using the HGG/LGG classifier

**Figure 3:**
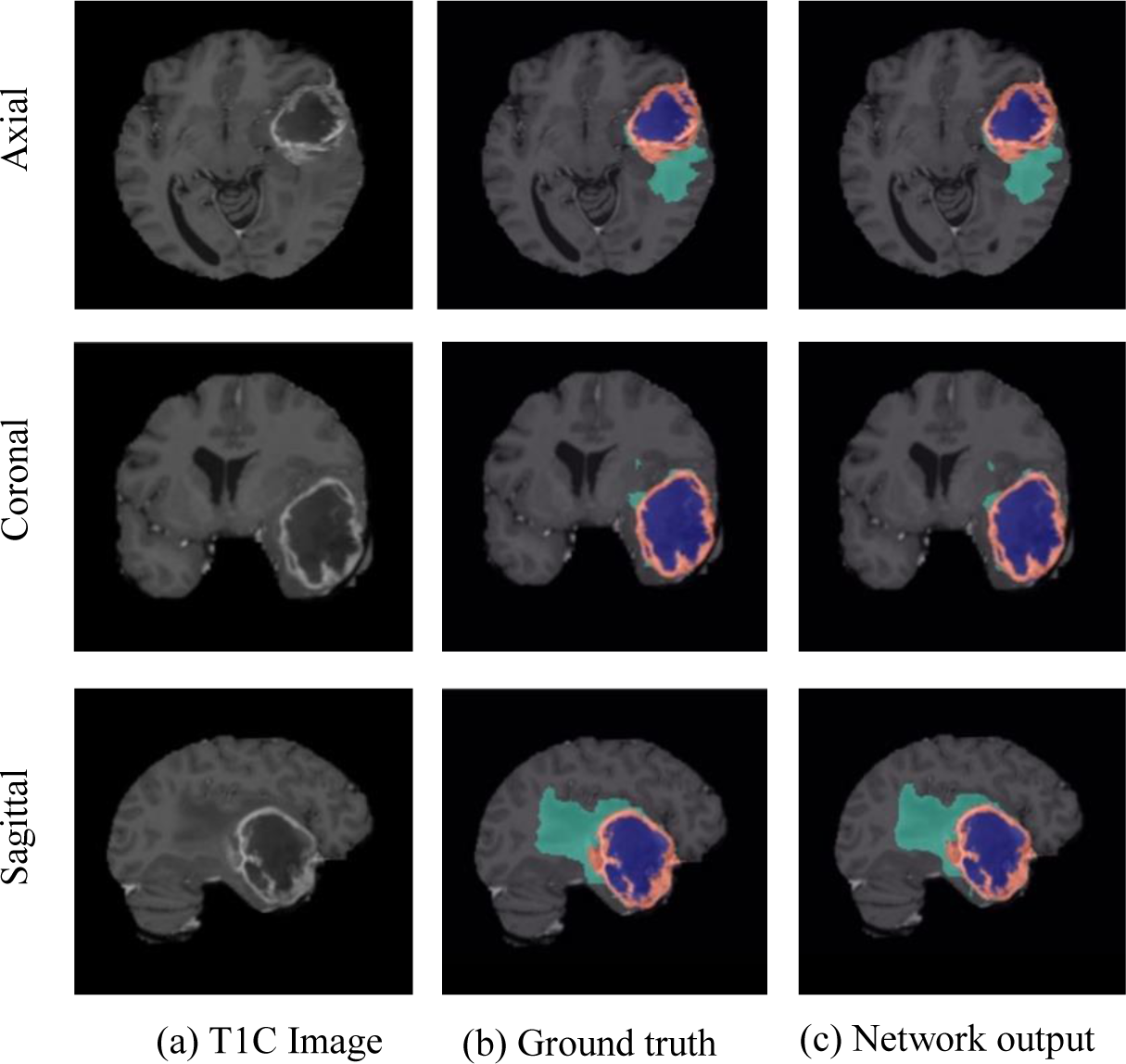
Example segmentation result for a High Grade Glioma (HGG) (a) A post contrast image. (b) Ground truth (c) Network output. Color Code: Red = Enhancing tumor, Blue = tumor core (Enhancing tumor + non-enhancing tumor + necrosis), Green = Edema, Whole tumor = Green + Blue + Red.

**Figure 4:**
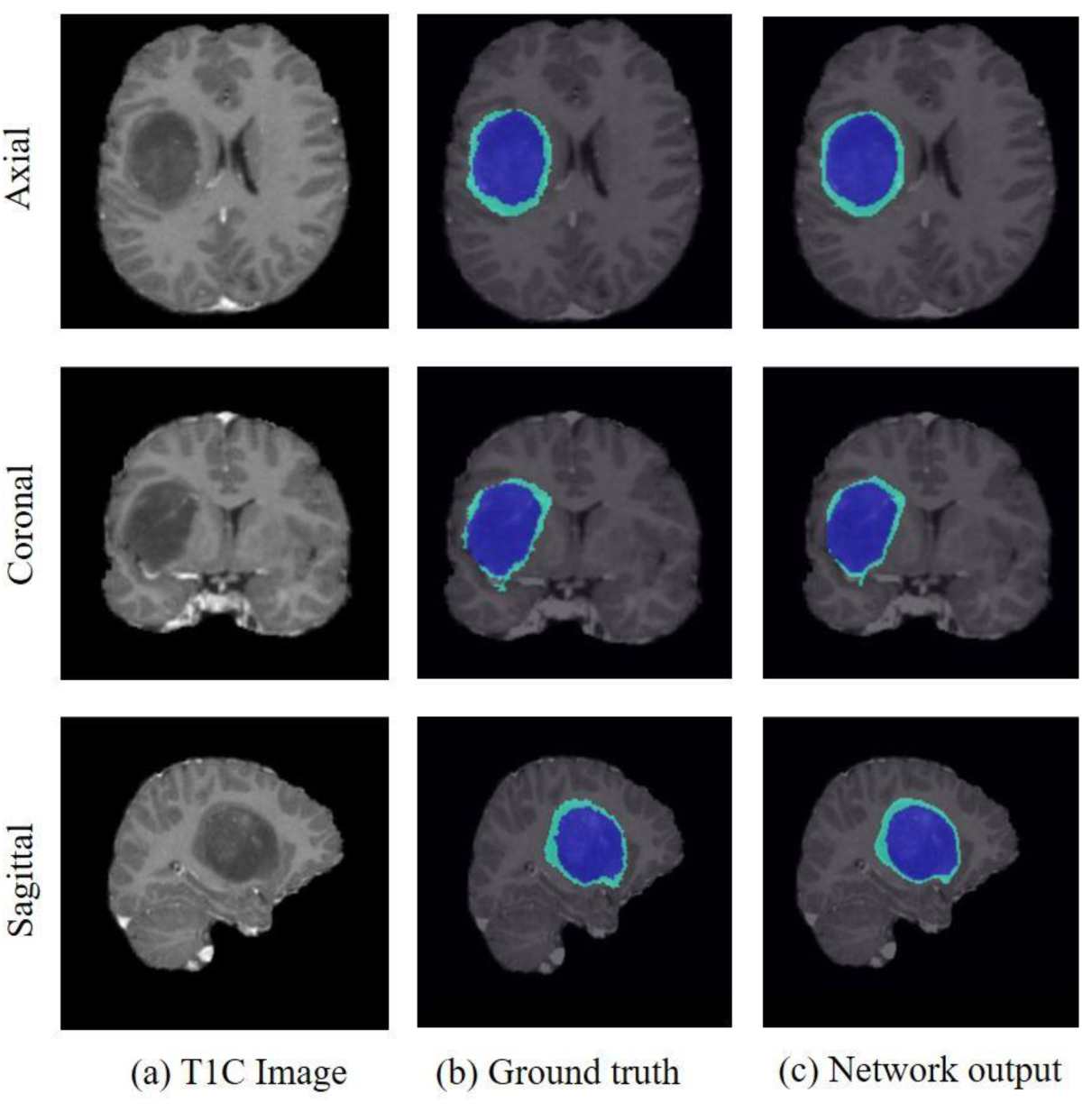
Example segmentation result for a Low Grade Glioma (HGG) (a) A post-contrast image. (b) Ground truth (c) Network output. Color Code: Red = Enhancing tumor, Blue = tumor core (Enhancing tumor + non-enhancing tumor + necrosis), Green = Edema, Whole tumor = Green + Blue + Red.

### 2.3 Training

#### 2.3.1 Brain Tumor Segmentation

Three groups of three Dense-UNets were designed. Each group consisted of networks designed to segment whole tumor, tumor core and enhancing tumor. Group 1 was trained using the dice-coefficient [27] as the loss function while group 2 was trained with binary focal loss [28] as the loss function. Group 3 consisted of 2 sub-groups namely, HGG group and LGG group. As the first step of group 3, a simple convolutional neural network was developed to separate HGG from LGG. The networks in the HGG group were trained using HGG cases only and the networks in the LGG group used the LGG cases only.

All 335 cases from the BraTS2019 dataset were used for training the networks from group 1 and group 2. 259 HGG cases were used to train the networks from the HGG group and 76 LGG cases were used to train the networks from the LGG group. 75% overlapping patches were extracted from the multi-parametric brain MR images that had at least one non-zero pixel on the corresponding ground truth patch. 20% of the extracted patches were used for in-training validation. Data augmentation steps included horizontal flipping, vertical flipping, random rotation, and translational rotation. Down sampled data (128×128×128) was also provided in the training as an additional data augmentation step. To circumvent the problem of data leakage, no patch from the same subject was mixed between training and in-training validation [29, 30].

#### Dice loss

The Dice co-efficient determines the amount of spatial overlap between the ground truth segmentation (X) and the network segmentation (Y)

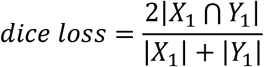

#### Focal loss

The focal loss was designed to address the problem of class imbalance between the foreground and background classes during training.

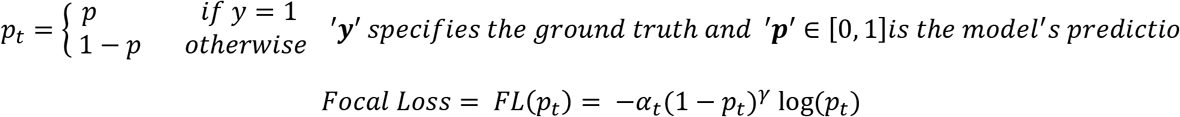

#### HGG & LGG Classifier

335 cases from the BraTS2019 dataset including 259 HGG cases and 76 LGG cases were used for training the network. Data augmentation steps included horizontal flipping, vertical flipping, random and translational rotation. The dataset was randomly shuffled and split into 60% (155 HGG and 46 LGG) for training, 20% (52 HGG and 15 LGG) for in-training validation and 20% (52 HGG and 15 LGG) for testing.

#### 2.3.3 Survival Analysis

Pyradiomics (a python package) was used to extract imaging (or radiomics) features [31]. Multi contrast MR images along with ground truth labels for each of the subcomponents were used to extract features from the BraTS2019 training dataset. For the BraTS2019 validation dataset, the segmented labels from the three 3D Dense UNets were used to extract features from the multi contrast MR images. 106 features were extracted for each MR imaging sequence from the tumor subcomponents (enhancing, edema and non-enhancing tumor with necrosis). Using these combinations, a total of 1272 features were extracted. Pywavelet, a python toolbox, was also used to extract wavelet based features [32]. 8 wavelet components for each MR image were extracted from level 1 of wavelet transform using coiflets (order=1) [33]. These 8 components (extracted from pywavelet toolbox) for the four MR imaging sequences in combination with 3 tumor subcomponents were used to extract 10,176 features using the pyradiomics package.

These imaging based features were combined with additional features including surface area, volume of tumor core, volume ofwhole tumor, ratio of tumor subcomponents to tumor core, ratio of tumor subcomponents to tumor volume, ratio of tumor core to tumor volume, and variance of enhancing tumor with T1C. The degree of angiogenesis was calculated by subtracting T1w and T1C in the tumor ROI, followed by a threshold of 50%. Finally, age and resection status were added to the feature set [34]. A total of 11,468 features were extracted combining the above features including imaging, texture and wavelet based features. Feature reduction was used to reduce the large number of features based on the feature importance determined by the gradient boost model. This reduced the feature space from 11,468 to 17 features. Table 1 shows a list of the selected 17 features for survival analysis. The 17 features were a combination of 10 Imaging features, 6 wavelet-imaging features and one non-imaging feature. All networks were trained using Tensorflow [35] backend engines, the Keras [36] python package and Pycharm IDEs on Tesla V100s and/or P40 NVIDIA-GPUs.

**Table 1:** Selected 17 features for survival analysis

| Feature No. | MR Imaging sequence | Tumor Mask | Pyradiomics Feature Name | Wavelet Component | Feature Category |
| --- | --- | --- | --- | --- | --- |
| 1 | T1C | Necrosis | original_glszm_SizeZoneNonUniformity | NA | Imaging |
| 2 | T2w-FLAIR | Enhancing | original_firstorder_Skewness | NA | Imaging |
| 3 | T2w-FLAIR | Necrosis | original_glszm_LargeAreaLowGrayLevelEmphasis | NA | Imaging |
| 4 | T2w-FLAIR | Edema | original_glszm_SmallAreaLowGrayLevelEmphasis | NA | Imaging |
| 5 | T2w | Enhancing | original_firstorder_Kurtosis | NA | Imaging |
| 6 | T2w | Enhancing | original_firstorder_Maximum | NA | Imaging |
| 7 | T2w | Necrosis | original_shape_Maximum2D DiameterRow | NA | Imaging |
| 8 | T1w | Enhancing | original_firstorder_Range | NA | Imaging |
| 9 | T1w | Enhancing | original_firstorder_Skewness | NA | Imaging |
| 10 | T1w | Edema | original_firstorder_Minimum | NA | Imaging |
| 11 | T2w-FLAIR | Enhancing | original_firstorder_Skewness | Component 1 | Wavelet- Imaging |
| 12 | T1C | Necrosis | original_glszm_HighGrayLevelZoneEmphasis | Component 2 | Wavelet- Imaging |
| 13 | T1C | Edema | original_firstorder_Minimum | Component 2 | Wavelet- Imaging |
| 14 | T1C | Edema | original_glszm_GrayLevelNonUniformity | Component 4 | Wavelet- Imaging |
| 15 | T1C | Edema | original_firstorder_Minimum | Component 4 | Wavelet- Imaging |
| 16 | T1w | Edema | original_glszm_SizeZoneNonUniformity | Component 5 | Wavelet- Imaging |
| 17 | Age | NA | NA | NA | Non Imaging Feature |

**Table 2:**
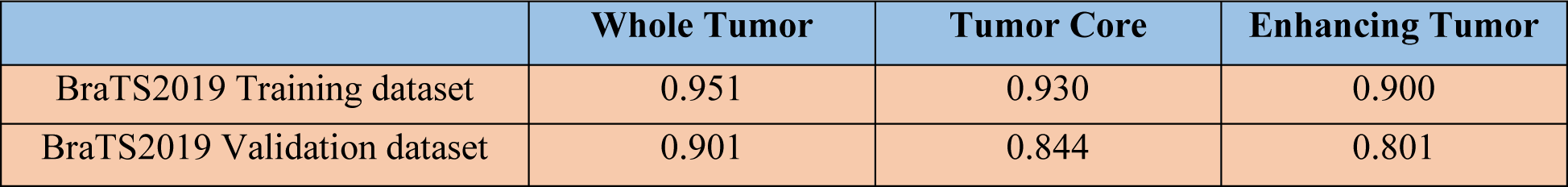
Mean dice scores on BraTS2019 datasets

**Table 3:** Uncertainty task results

|  | Dice AUC<br>Whole<br>tumor | Dice AUC<br>Tumor<br>core | Dice AUC<br>Enhancing<br>tumor | FTP Ratio<br>AUC Whole<br>tumor | FTP Ratio<br>AUC<br>Tumor core | FTP Ratio<br>AUC<br>Enhancing |
| --- | --- | --- | --- | --- | --- | --- |
| BraTS2019<br>Validation dataset | 0.894 | 0.887 | 0.876 | 0.012 | 0.061 | 0.046 |

### 2.4 Testing

#### 2.4.1 HGG & LGG classifier

The classifier was evaluated on 67 cases including 52 HGGs and 15 LGGs. To generalize the network’s performance, a 3 fold cross-validation was also performed. While we used the HGG and LGG classifier as the first step for brain tumor segmentation, the BraTS2019 validation dataset did not include labels for HGG and LGG precluding evaluation of classification accuracy for this initial step on the validation dataset.

#### 2.4.2 Brain Tumor Segmentation

All the networks were tested on 125 cases from the BraTS2019 validation dataset. Patches of size 32×32×32 were provided to the networks for testing. The prediction patches were then used to reconstruct a full segmentation volume. Each group was tested in 3 different ways including (a) non-overlapping patches, (b) 25% overlapping patches and (c) 50% overlapping patches. At the end of testing, each group produced 3 segmentation volumes for a particular label resulting in 9 segmentation volumes across the 3 groups. The 9 segmentation volumes were assigned with equal weights, averaged and thresholded at 0.5 for enhancing tumor, 0.8 for whole tumor and 0.7 for tumor core for every voxel. The same procedure was performed on whole tumor, tumor core and enhancing tumor labels. The ensembled output of whole tumor, tumor core and enhancing tumor were fused in a post-processing step that included the 3D connected components algorithm to improve prediction accuracy by removing false positives.

#### 2.4.3 Survival Analysis

The network was evaluated on 29 cases from the BraTS2019 validation dataset. The Survival analysis was evaluated only for the GTR cases. The predicted overall survival (OS) task classified the subjects as long-survivors (greater than 15 months), mid-survivors (between 10 to 15 months) and short-survivors (less than 10 months).

#### 2.4.3 Uncertainty task

Uncertainty masks were also created for the 125 cases from the BraTS2019 validation dataset. The predicted probability maps for a particular label from each group were assigned with equal weights, averaged and scaled from 0 to 100 such that 0 represents the most certain prediction and 100 represents the most uncertain. In this task, uncertain voxels were removed at multiple predetermined threshold points. The performance of the networks was assessed based on the dice score of the remaining voxels.

## 3. RESULTS

### 3.1 HGG & LGG Classifier

The network achieved a testing accuracy of 89% and an average cross-validation accuracy of 90%.

### 3.2 Brain Tumor Segmentation

This method achieved average dice scores of 0.95, 0.93 and 0.90 on WT, TC and ET on the training dataset (Table 1). It also achieved average Dice scores of 0.901, 0.844, 0.801 and sensitivity of 0.924, 0.846, 0.796 specificity of 0.991, 0.999, 0.998 for WT, TC and ET respectively on the 125 held-out cases along with Hausdorff distances of 7.60, 8.31 and 5.49 mm respectively.

### 3.3 Survival task

The linear regression model achieved an accuracy of 55.2% and mean squared error (MSE) of 119244.6 (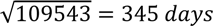) with median SE of 19881 and standard deviation of 227388.7 and spearman correlation co-efficient of 0.271.

### 3.4 Uncertainty task

The networks achieved average dice_AUC of 0.894, 0.887, 0.876 and FTP_AUC_ratio of 0.012, 0.061 and 0.046 for WT, TC and ET respectively on the BraTS2019 validation dataset.

## 4. DISCUSSION

Accurate, efficient, and reliable tumor segmentation algorithms have the potential to improve management of GBM patients. Currently the vast majority of clinical and research efforts to evaluate response to therapy utilize gross geometric measurements. MRI-based glioma segmentation algorithms represent a method to reduce subjectivity and provide accurate quantitative analysis to assist in clinical decision making and improve patient outcomes. Although the underlying task can be simply stated as a voxel-level classification, a wide variety of automated and semi-automated tumor segmentation algorithms have been proposed. In this work, we developed a fully automated deep learning method to classify gliomas as high grade and low-grade, segment brain tumors into subcomponents and predict overall survival. This method was tested on the BraTS2019 validation dataset including 125 cases for the tumor segmentation task and 29 cases for the survival analysis task. A three group framework was used, providing several advantages compared to the currently existing methods. Using three binary segmentation networks for segmenting the tumor into its sub-components allowed us to use a simpler network for each task [14]. The networks were easier to train with reduced over-fitting [14]. Furthermore, since all three networks were trained separately as binary segmentation problems, misclassification was greatly reduced, as demonstrated by the results from the uncertainty task.

## 5. CONCLUSION

Fully automated convolutional neural networks were developed for segmenting brain tumors into their subcomponents. This algorithm reached high performance accuracy on the BraTS2019 validation dataset. High Dice scores, accuracy and speed of this network allows for large scale application in brain tumor segmentation. This method can be implemented in the clinical workflow for reliable tumor segmentation, survival prediction and for providing clinical guidance in diagnosis, surgical planning and follow up assessments.

